# OsbZIP47 an integrator for meristem regulators during rice plant growth and development

**DOI:** 10.1101/2021.05.11.443658

**Authors:** Sandhan Prakash, Rashmi Rai, Raghavaram Peesapati, Usha Vijayraghavan

## Abstract

Stem cell homeostasis by the WUS-CLV negative feedback loop is generally conserved across species; however, its links with other meristem regulators may have species-specific distinctions, rice being an example. We characterize rice OsbZIP47 for vegetative and inflorescence phenotypes in knockdown (OsbZIP47KD) transgenics and uncover its role in meristem maintenance and developmental progression. The shoot apical meristem (SAM) size in five day old OsbZIP47KD seedlings, was reduced as compared to the wild-type (WT). Whereas SAM in older twenty-five-day OsbZIP47KD plants was larger with increased size for L1 and underlying cells. We tested protein interactions of OsbZIP47 with other transcription factors and found partnerships with OsMADS1, RFL, and OSH1. Results from meta-analysis of deregulated panicle transcriptome datasets, in OsbZIP47KD, OsMADS1KD and RFLKD knockdown transgenics, and OSH1 genome-wide binding sites divulge potential targets coregulated by OsbZIP47, OsMADS1, OSH1 and RFL. Transcript analysis in OsbZIP47KD SAM and panicles showed abnormal gene expression for CLAVATA peptide-like signaling FON2-LIKE CLE PROTEIN1 (FCP1), FLORAL ORGAN NUMBER 2 (FON2), and hormone pathway: cytokinin (CK) Isopenteyltransferase2 (OsIPT2), Isopenteyltransferase8 (OsIPT8); auxin biosynthesis OsYUCCA6, OsYUCCA7; gibberellic acid (GA) biosynthesis GA20Ox1, GA20Ox4 and brassinosteriod biosynthesis CYP734A4 genes. The effects on ABBERANT PANICLE ORGANIZATION1 (APO1), OsMADS16, and DROOPING LEAF relate to second and third whorl organ phenotypes in OsbZIP47KD florets. Further, we demonstrate that OsbZIP47 redox status affects its DNA binding to cis elements in the FCP1 locus. Taken together, we provide insights on unique functional roles for OsbZIP47 in rice shoot meristem maintenance, its progression through inflorescence branching and floret development.

**One sentence summary:** OsbZIP47 regulates rice shoot meristem size, panicle and floret development in concert with other meristem regulators such as OsMADS1, RFL and OSH1.

## INTRODUCTION

The post-embryonic development in flowering plants depends on the balance between stem cell renewal in the central zone of above ground meristems and the adoption of specific differentiation programs from the peripheral zone. The genetic framework of the basic WUS-CLV pathway for meristem maintenance is largely conserved in monocots and dicots yet some functional differences are reported among the cereal grass models maize and rice. The tissue-specific effects of a pair of paralogous rice *CLV3-like* genes, that encode the ligand in this signalling pathway (Suzaki et al., 2004; Suzaki et al., 2006), and the male *vs.* female inflorescence specific roles for maize *CLV1* and *CLV2* homologs, *THICK TASSEL DWARF1* (*TD1*) and *FASCIATED EAR2* (*FEA2)*, exemplify species-specific innovations in components of this core meristem regulatory circuit (reviewed in Bommert et al., 2005b; Dodsworth 2009; Paulter et al., 2013; Chongloi et al., 2019). *FLORAL ORGAN NUMBER 1* (*FON1*) is the rice ortholog of *CLV1*, while *FON2/FON4*, *FON2 SPARE1* (*FOS1*) and *FCP1* (*FON2-LIKE CLE PROTEIN1*) are CLV3 peptide paralogs. Interestingly, FON2 signaling through FON1 majorly regulates homeostasis in the inflorescence meristems whereas FCP1 triggered signaling regulates the vegetative shoot apical meristem (SAM) through effects on *OsWOX4*; functionally related to *AtWUS* (Nagasawa et al., 1996, Suzaki et al., 2004, 2006; Ohmori et al., 2013). Integration of CLV-WUS pathway with the roles of class I *Knotted-1-like homeobox* (*KNOX*) genes, *Arabidopsis STM*, rice *OSH1* and maize *KNOTTED1* (*KN1*) in meristem maintenance is conserved across species (Long et al., 1996; Tsuda et al., 2011; Vollbrecht et al., 2000). Similarly, the interlinking of WUS-CLV pathway with phytohormone-based meristem control by cytokinin (CK), auxin (IAA/AUX), Gibberellin (GA), Brassinosteriod (BR) is also conserved (Lee et al., 2009; Gordon et al., 2009; Kurakawa et al., 2007; Yamaki et al., 2011; Zhao et al., 2010, Somssich et al., 2016).). The feed-forward regulatory loop of *KNOXI* homeobox transcription factor genes, *AtSTM*/*ZmKN1*/*OSH1*, maintains the SAM by crosstalk with cytokinin (CK) pathway and *AtWUS* also regulates cytokinin signaling (Kerstetter et al., 1997; Brand et al., 2002; Leifried et al., 2005; Bolduc and Hake, 2009; Tsuda et al., 2011, 2014, Ohmori et al., 2013; Gordon et al., 2009; Kurakawa et al., 2007). Despite several conserved components for meristem regulation among eudicot and monocot species, the interconnecting regulators are poorly understood in cereals.

In the floral meristem center the timing of the termination of stem cell activity is co-incident with carpel/ovule specification. This creates a determinate floral meristem for normal reproduction. Meristem termination is mediated by the concerted activity of floral organ identity genes (Class A, C and E) whose regulatory effects on *WU*S-*CLV* and/or *WUS*-*KNOX* pathway genes operates in both monocots and eudicots (reviewed in Tanaka et al., 2013; Callens et al., 2018; Chongloi et al., 2019). These homeotic genes are in turn spatially and temporally regulated. For example, *AtLEAFY* directly activates *AP1*, and *WUS* (Lenhard et al., 2001; Lohmann et al., 2001) and repress shoots meristem identity gene *TERMINAL FLOWER1* (*TFL1*) (Moyroud et al., 2009, 2010). Furthermore, in young floral meristems LEAFY (LFY) together with UNUSUAL FLORAL ORGAN (UFO) and WUS activates *AP3* and *AG* gene expression in third and fourth whorls of the developing meristem (Parcy et al., 1998; Busch et al., 1999; Wagner et al., 1999; Lenhard et al., 2001; Lohmann et al., 2001). In later developmental stages the activation of *KNUCKLES* (*KNU*), by AG, leads to repression of *WUS* by the recruitment of Polycomb group (PcG) chromatin modifiers (Ming and Ma, 2009; Sun et al., 2009, 2014; Liu et al., 2011; Zhang, 2014). Aside from LFY, other regulators of *AG* expression are known, one of which is *PERIANTHIA* (*PAN*) that encodes a bZIP class TF, whose orthologs are *OsbZIP47* and *ZmFEA4* in rice and maize respectively. Floral meristem size and organ patterning defects in Arabidopsis *pan-3 lfy-31* double mutant, and in transgenics with modified PAN fusion proteins (repressive *vs*. activated forms) show its role in floral determinacy, meristem size and floral organ patterning (Chuang et al.,1999; Running and Meyerowitz, 1996; Das et al., 2009; Maier et al., 2009, 2011). Maize *ZmFEA4* (*FASCIATED EAR4*) is a positive regulator of lateral organ differentiation in both vegetative and reproductive growth phases. Its effects on auxin pathway, balances *WUS* and *KNOTTED1* (*KN1*) based meristem homeostasis and promotes the expression of lateral organ genes *YABBY3*, *KANADI*, *ASYMMETRIC LEAVES2*, *CUC* among several others (Pautler et al., 2015). Functions for its rice homolog, *OsbZIP47* (LOC_Os06g15480) are not well studied, however extensive genetic and molecular studies have identified many other rice meristem homeostasis factors. For instance during rice inflorescence (panicle) development ABERRANT PANICLE ORGANIZATION 1 (APO1), ortholog of AtUFO and APO2/RFL, ortholog of LFY, interact and promote panicle branch meristem (BM) indeterminacy thereby regulate spikelet meristem (SM) to floral meristem (FM) transition (Ikeda et al., 2005, 2007; Rao et al., 2008, Ikeda-Kawakastu et al., 2009, 2012). Other rice inflorescence branch meristem identity and transition regulators are closely-related gene members of the AP2-ERF family: *OsIDS* (*INDETERMINATE SPIKELET*), *SNB* (*SUPERNUMERARY BRACT*), *FZP* (*FRIZZY PANICLE*), *MFS1* (*MULTIPLE FLORET SPIKELET1*). These genes control flowering time and progression through panicle branching by regulating other classes of genes like *RFL* and the MADS-box genes, *OsMADS1* and *OsMADS6*, among others (Lee and An, 2012; Komatsu et al., 2003, Ren et al., 2013). How these inflorescence BM identity and transition regulators intersect with the core canonical CLV-WUS and WUS-KNOXI meristem maintenance circuits is not much explored in rice or maize. Our studies on phenotypic and functional characterization of rice *OsbZIP47* by RNA interference (dsRNAi) based knockdown (KD) give clues on its role in meristem size and identity, and on downstream pathways that can be co-regulated by OsbZIP47 and OSH1, OsMADS1 and RFL.

## RESULTS

### *OsbZIP47* knockdown plants have abnormal SAM Size

To investigate developmental roles of OsbZIP47 we generated *OsbZIP47* knockdown transgenics (*OsbZIP47*KD) by RNA interference (dsRNAi). Phenotypic analysis in the T3 generation for one of the lines: *OsbZIP47K*D line #14 was done. In pooled panicle tissues (0.1-0.5cm) from this line qRT-PCR showed ∼24-fold down regulation of the endogenous *OsbZIP47* transcripts (Supplemental Fig. S1). The earliest phenotype noted was the seedling height at 8 days after germination (DAG), which was significantly reduced in *OsbZIP47KD* as compared to wild type (*WT*) (Fig. 1B and C, Supplemental Table S1). This observation lead us to investigate the size of shoot apical meristem (SAM) in histological sections of 5 DAG and 25 DAG seedlings from both wild type (*WT*) and *OsbZIP47KD* plants. Interestingly, the 5 day old *OsbZIP47KD* seedlings show reduced SAM (895.7 μm^2^) area as compared to that in *WT* (1131 μm^2^); whereas in 25-day *OsbZIP47KD* seedlings SAM area was contrastingly increased (Fig. 1E). To understand the cellular differences that underlie these meristem abnormalities, the number of cells in L1 layer, were counted to identify reduced number of L1 cells in 5 and 25 DAG *OsbZIP47 KD* seedlings (Supplemental Fig S2). Further, L1 layer cell size in apical and basal regions (the peripheral zone) of the meristem and for internal cells underlying the L1 layer (as marked in Fig.1D) was measured. In 5 DAG seedlings we observed reduced L1 cell size in the apical region whereas L1 cell size in the basal/ peripheral and cell size in internal regions of meristem were not altered (Fig. 1F). This suggests that the compromised overall SAM area in 5 DAG seedling is probably due to decrease in the number and size of apical L1 cells. Intriguingly, in 25 DAG *OsbZIP47KD* seedlings L1 cell size in apical and basal/peripheral regions was increased. Strikingly, cells underlying the L1 apical region were larger (Fig. 1D, F). This suggests the increased SAM area in 25 DAG seedlings is probably not due to increased cell number but rather due to enlarged cell size. The spatial distribution of dividing cells was assessed by RNA *in-situ* for the cell cycle S-phase maker HISTONE4 (H4) (Supplemental Fig. S3). Compared to the *WT*, the overall signal in *OsbZIP47KD* tissues (5 D and 25D) was higher. Also, we observed unexpected higher signals in the meristem central region, mainly in 25 DAG seedling which was reminiscent to the pattern in *sho* mutant in *OSH1* (Itoh et al., 2000). Together these data on likely altered cell division patterns in the SAM of *OsbZIP47KD* plants (Supplemental Fig. S3) indicate a regulatory role for OsbZIP47 in meristem (SAM) maintenance. To understand the molecular signatures relating to these meristem defects we tested transcript levels for few known SAM regulators (Fig. 1G) in 25 DAG seedlings. The down-regulation of *FCP1* and *FON2* (rice homologs of *CLV3*), *APO1* (*UFO1* homolog), *CYP734A4* and *YUCCA6* was observed. *CUC1,* lateral meristem boundary marker, showed a marginal reduction in expression. These gene expression effects of OsbZIP47 can be related to abnormal SAM maintenance on its knockdown with novel effects on components in the CLV-WUS pathway, on other meristem regulators, auxin and brassinosteriod phytohormone pathways.

**Figure 1.**
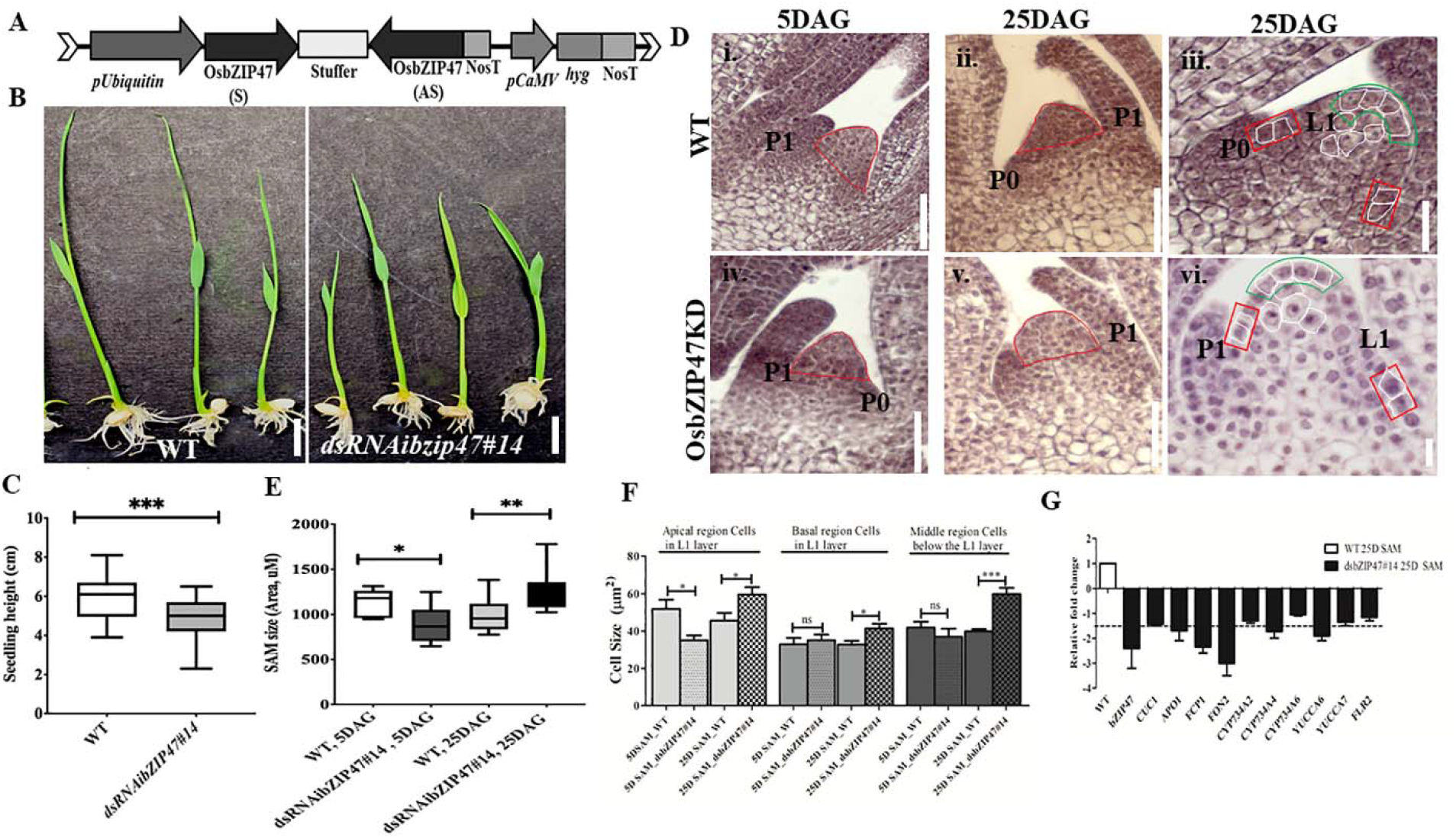
*OsbZIP47* knockdown (KD) vegetative phenotypes. (A) Schematic representation of *dsRNAiOsbZIP47* transgene T-DNA segment. (B) Seedling height of 8 day *OsbZIP47KD* plants is shorter than that of wild type (*WT*) plants. Scale bar =1 cm. (C) Seedling plant height data shown as mean ± s.d., Student’s *t* tests, ***P< 0.0001, n=30. (D) and (E) Comparison of SAM area in *OsbZIP47KD vs. WT* in 5 DAG (days after germination) and 25 DAG seedlings. SAM area, marked by red outline in panels i, ii, iv, and v, is reduced in 5 DAG *OsbZIP47KD* seedlings but is increased in 25 DAG seedlings. Scale bar in i, ii, iv, and v are 50 mM, and iii and vi are 20 mM. Data is shown as mean ± s.d. Student’s *t* tests, *P< 0.01, **P< 0.001, n=10. (F) Comparison of cell size in the apical region, the internal underlying apical L1and in the peripheral zone of the meristem. The average of 5 cells was taken as a data point for a single section and 10-12 different sections were taken for statistical analysis. Asterisks indicate significant differences from wild type at * P<0.01, in 5 DAG SAM, *** P<0.0003 in 25 DAG SAM tissues respectively. NS denotes non-significant difference. (G) RT-qPCR analyses of *CUC1, APO1, FCP1, FON2, CYP734A2, CYP734A4, CYP734A6, YUCCA6, FLR2* and FON2 transcripts from candidate targets in SAM from 25 DAG seedlings. Fold change values were determined by comparing the normalized expression levels in *OsbZIP47KD* plants to *WT* plants.

### Late heading date and altered panicle architecture on *OsbZIP47* knockdown

*OsbZIP47KD* plants are delayed by 20 days for flowering i.e., SAM to inflorescence meristem (IM) transition. At this stage *OsbZIP47KD* plant height was reduced as compared to *WT* (Fig. 2A to D) and suggests that OsbZIP47 promotes the transition from vegetative to the reproductive phase. The shorter plant height was due to poor stem internode elongation in the KD transgenics without change in internode number (Fig. 2F). The panicle of knockdown plants showed developmental abnormalities: reduced inflorescence axis (panicle rachis) length, lowered primary branch (Fig. 2E, and spikelet number (Fig. 3 A, L and M; Supplemental Table 1). These events together indicate an early progression of primary branch meristems to spikelets in these *OsbZIP47KD* plants and point to a possible role of OsbZIP47 in the temporal control of branch meristem indeterminacy. Moreover, *OsbZIP47KD* plants had greater flag leaf lamina joint angle as compared to the *WT* (Fig. 3B, C, K). Mutants in *cpy734a4* (Tsuda et al., 2014), a downstream gene target of OsbZIP47 (see Fig. 1G) also share this phenotype and have abnormal meristems. Plant architecture, flowering time, leaf angle and inflorescence architectures all impact yield, grain shape and size (Harder and Prusinkiewicz, 2013; Sakuma and Schnurbusch., 2019). Interestingly the seeds from *OsbZIP47KD* plants were altered for the length/width (L/W) ratio as compared to WT seeds (Fig. 3I and J).

**Figure 2.**
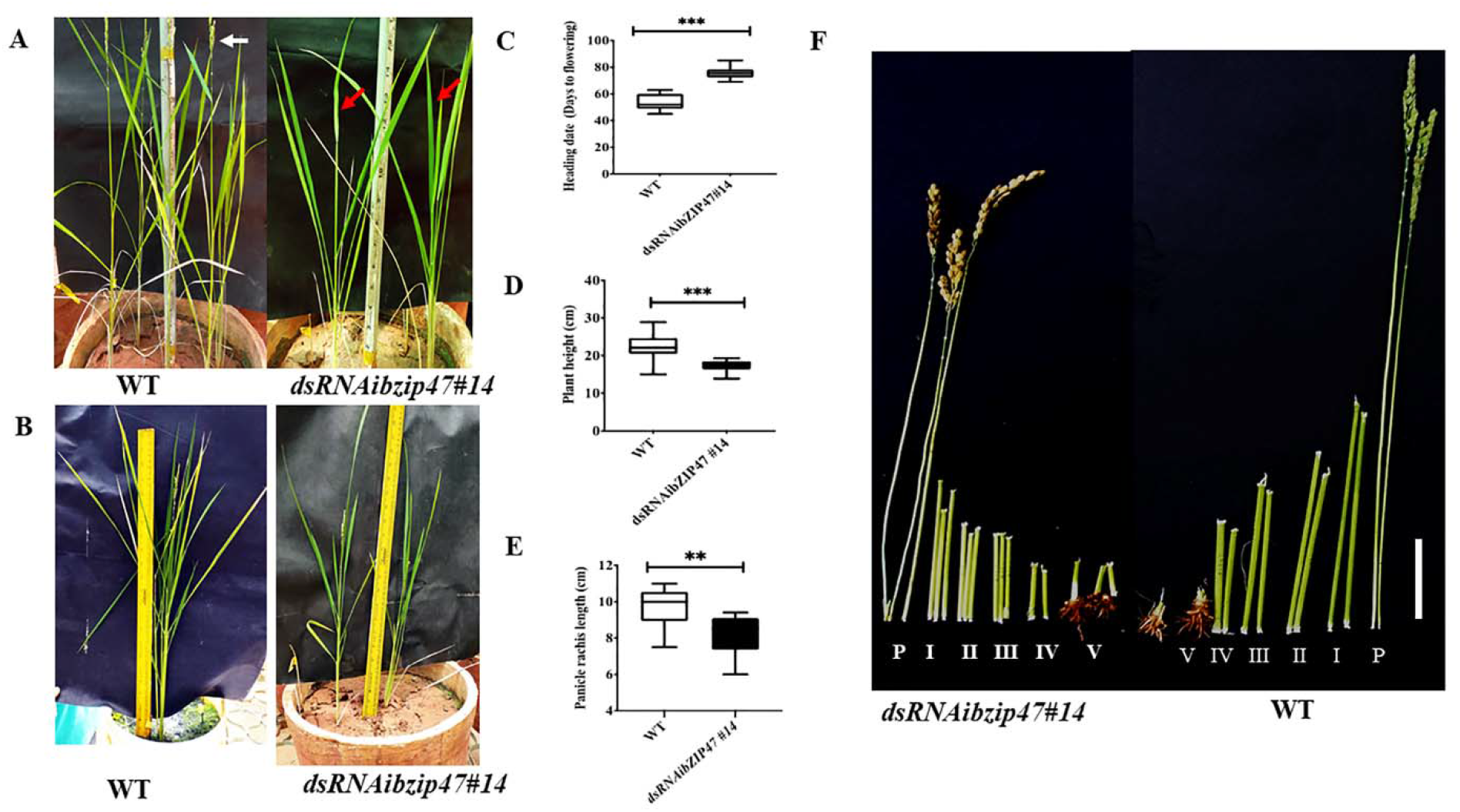
Phenotypic effects of *OsbZIP47KD* on plant growth and floral transition. (A) Flowering timing (SAM to Inflorescence Meristem/ panicle meristem transition) in *OsbZIP47KD* plants is delayed by ∼22days as compared to the *WT*. White arrow indicates the booted panicle in *WT* and red arrow points the absence of the booted panicle in a *OsbZIP47KD* plant of the same age. (B) Plant height in mature flowering plants shows as compared to the WT, *OsbZIP47KD* plants are shorter. Quantitation of heading dates (C) plant height (D) and panicle rachis length (E) in *WT* and *OsbZIP47KD* plants. Data are shown as the mean ± s.d. (Student’s t tests, **P < 0.01, n = 10). (F) Internodes in mature flowering *WT* and *OsbZIP47KD* plants are displayed and are numbered I to V from the apical end. P is the panicle bearing node and internode. Bar, 2.5 cm. Shorter internodes I to V in *OsbZIP47KD* contributes to the reduced height.

**Figure 3.**
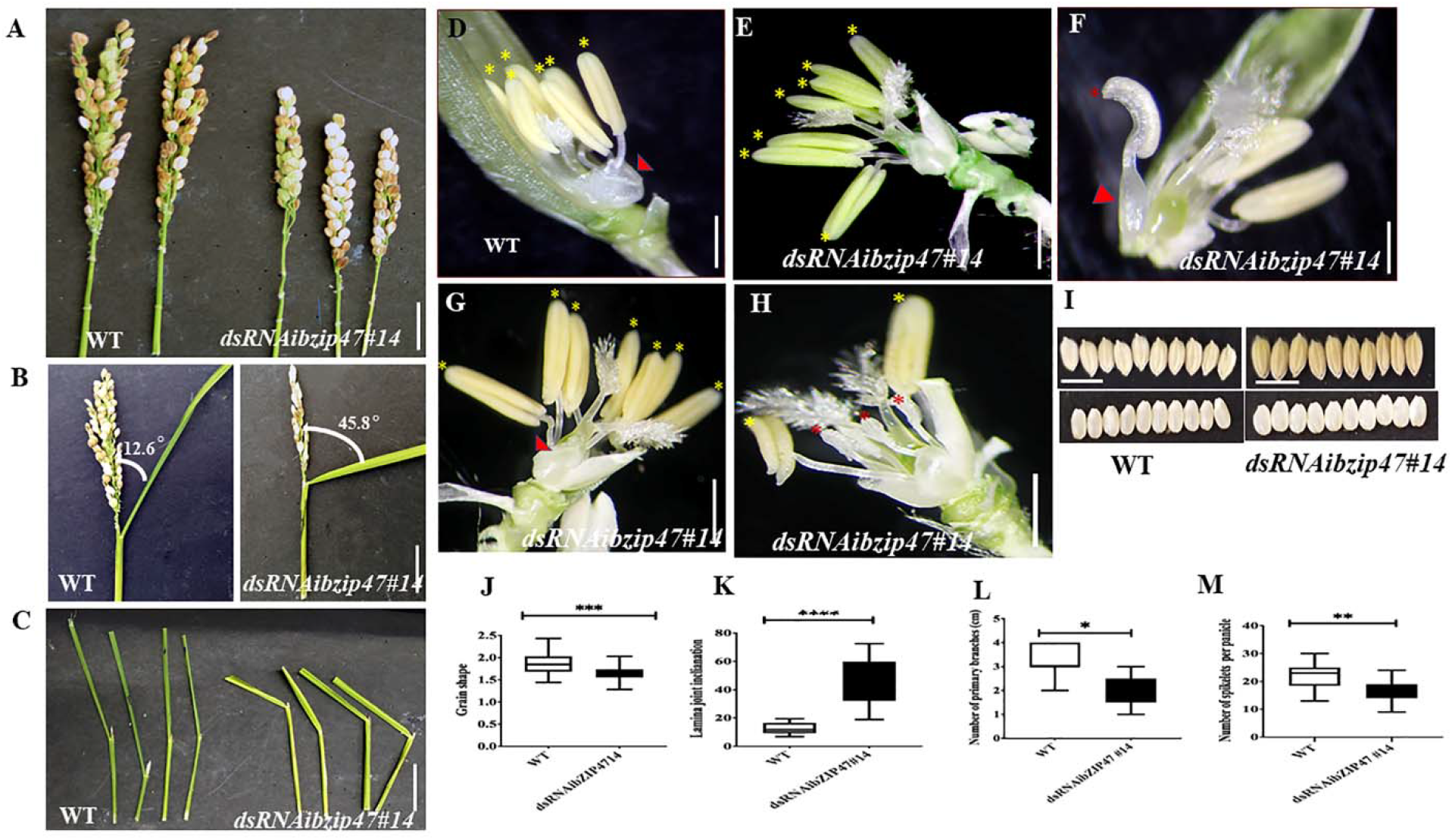
Floret organ numbers and organ development in *OsbZIP47KD* plants. (A) Shorter panicle *OsbZIP47KD* with reduced number of primary branches and spikelet number per panicle as compared to wild type plant. (B) and (C), Flag leaf angle in fully headed panicles of mature plants show increased lamina joint angle in*OsbZIP47KD* plant as compared to the *WT*. (D) *WT* floret organs shown after removal of lemma and the spikelet sterile lemmas. Red arrow points to the pair of lodicules and yellow asterisks to the six normal stamens. (E-H), Organ phenotypes in *OsbZIP4*7KD florets. (E) Floret with 7 stamens (yellow asterisks) with normal filaments and anthers. (F) Floret with mildly deformed lodicule (red arrowhead), two of the six normal stamens (others were dissected out) and a chimeric second whorl organ with lodicule and stamenoid tissues (red asterisk). (G) Floret with deformed lodicule and 7 normal stamens (yellow asterisk). (H) Floret with slightly deformed lodicule, short stamens and shrunken anthers (red asterisk), two near normal stamens (yellow asterisk). (I) and (J), Grain morphology in *OsbZIP47KD* and *WT* plants. The length/width (L/W) ratio of *OsbZIP47KD* seeds was lower than *WT* seeds, suggesting increased grain width. Scale bar =1cm in panels in A to I. (J-M), Statistical analysis (mean ± s. d.) of grain shape, lamina joint angle, primary branch number, number of spikelets per panicles. Data in J (n=40 in J), K (n=10), L (n=11), M (n=9), Student’s t tests, **P < 0.01, ***P<0.001.

### *OsbZIP47* contributes to second and third whorl, lodicule and stamen, organ development

*OsbZIP47KD* floret phenotype were largely restricted to lodicules, and stamens (Fig. 3D to 3H). The organ defects were grouped into four classes. In class I, 40% florets had mild deformed lodicule (distal elongation) with normal stamen number (Supplemental Fig. S4), class II ∼28% florets showed mild lodicule elongation with abnormal short stamens and indehiscent anthers (Fig. 3H). In florets of class III (∼20%) had partially deformed lodicules with an increase in stamen number to 7 (Fig. 3E and G). In the ∼12% class IV knockdown florets, mildly elongated lodicules co-occurred with chimeric organs usually with lodicule and stamen characteristics (Fig. 3F, G and H). Also, in most florets the lemma, was abnormally fused with lodicule making dissection of the lemma away from lodicule difficult. All together these data suggest *OsbZIP47* contributes to floral organ development in the second and third whorls. As a complementary analysis we examined consequences of ubiquitous overexpression of full-length cDNA of *OsbZIP47* in transgenic rice (Supplemental Fig. S5). None of the *OsbZIP47Ox* lines showed changes from the *WT* (data not shown). A speculation is that *OsbZIP47* functions may depend on partners or that some post-translational modifications (PTMs) may modulate its functions, as also concluded from overexpression studies of Arabidopsis *AtPAN* (Chuang et al., 1999).

### Tissue expression profile of OsbZIP47 through development

RNA *in situ* hybridization was performed to examine the spatial distribution of *OsbZIP47* mRNA. These experiments confirmed transcripts in various meristems, consistent with the phenotypes seen on *OsbZIP47KD*. Transcripts are evenly distributed in SAM of young seedlings (5 DAG) with slightly higher levels in the emerging leaf primordia (Fig. 4A). This pattern is somewhat different from maize *FEA4* where the signals are excluded from SAM stem cell niche and from incipient P0 leaf primordium (Paulter et al., 2015). The spatial pattern of *OsbZIP47* transcripts in SAM of 25 day plants is similar to that in 5 DAG plants (Fig. 4B). During reproductive development, high levels of *OsbZIP47* transcripts are at apical end of growing inflorescence meristem (IM/ rachis) and the ends of branch meristem (PBM and SBM) which relates to the poorly branched inflorescence of knockdown plants. In elongating primary and secondary branches (Fig. 4C and D), in spikelet meristem (SM, Sp2, Fig. 4E), and in floral meristems (Sp4-Sp6, Fig. 4F) the signal is spatially uniform. However, in mature florets *OsbZIP47* RNA was confined to the lodicule, stamen and carpel organ primordia and differentiating organs (Fig. 4G). Additionally, hybridization signal in carpel wall (c) and ovule (o) (Fig. 4H) was observed. Arabidopsis *pan* mutant flowers occasionally have multiple carpel up to three with deviated gynoecium (Running and Meyerowitz, 1996; Running et al., 1998). We speculate OsbZIP47 may have a minor role in carpel development or is functionally redundant fashion with floral C-class function genes. Thus, *OsbZIP47* is expressed in various above-ground meristems reflecting its diverse roles in different meristems.

**Figure 4.**
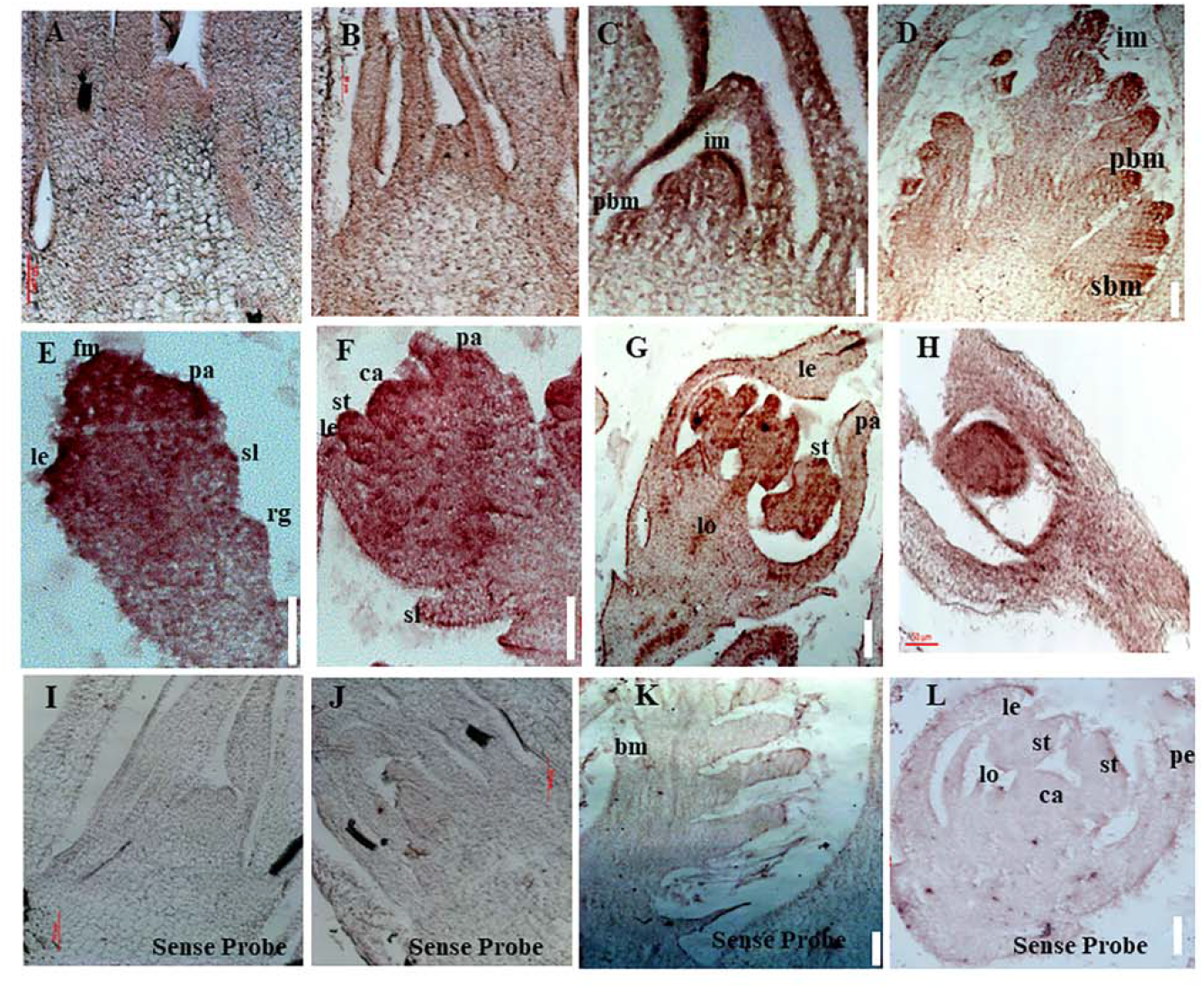
Spatial distribution of *OsbZIP47* transcripts in meristems and florets. (A and B) *OsbZIP47 in situ* RNA hybridization signal in SAM tissues from 5 DAG and 25 seedlings. Expression in SAM with slightly higher signal in emerging leaf primordia. (C and D) Inflorescence meristem (IM) with emerging primary branch meristems show expression of *OsbZIP47* in the elongation meristem and primary branch meristems (prb) and secondary branch meristems (srb) and in young leaves. (E) *OsbZIP47* transcripts in very early floret meristem (FM) with uniform spatial distribution of signal. (F) Floret uniform signal in the well-formed stamens (st), lemma/ palea (le and pa) organ primordia and in the central early carpel primordia. (G) High level *OsbZIP47* expression in the lodicule and in stamens, lower signal in the near mature lemma and palea. (H) *OsbZIP47* transcripts in the ovary wall and ovule. (I and J) SAM in 5 DAG and 25 DAG plants probe with sense probe as a negative control. (K and L) IM and near mature floret respectively probed with sense RNA.

### Heterodimerization of OsbZIP47 with other floral meristem regulators

Among the co-occurring motifs in genome-wide loci bound by OsMADS1, motif for bZIP factor binding is enriched (Khanday et al., 2016). This was the basis for our hypothesis that OsMADS1 and members of OsbZIP family could function in complexes in early floral meristems. Based on temporal co-expression profiles of *OsMADS1* and *OsbZIP47* (Arora et al., 2007; Nijhawan et al., 2008) we tested interaction among these proteins using the heterologous yeast two hybrid assay (Fig. 5). Additionally, to investigate possibility of OsbZIP47 heterodimerization with other meristem regulators, we relied on reports from genetic studies in Arabidopsis, or maize, or rice to curate and choose candidates (Das et al., 2009; Paulter et al., 2015; Khanday et al., 2013; Deshpande et al., 2015). OSH1, RFL, ETTIN1 and ETTIN2 emerged as candidates. We re-visited reports on the panicle and floret expression patterns of these meristem regulators to deduce if spatial co-expression of *OsbZIP47*, *OSH1* and *OSH15* could occur. RNA *in situ* patterns of *OSH1* in rice panicles and florets (Komatsu et al., 2001; Chu et al., 2006; Yoon et al., 2017) and *OsbZIP47* transcript spatial profile (Fig. 4) point to an overlap of *OsbZIP47* and *OSH1* transcripts in primary and secondary branch primordia and in a broad range of developing spikelet/ floret meristems (Sp2 to Sp8) (Supplemental Fig. S6). Moreover, Paulter et al., (2015) reported the several gene loci are bound by ZmKN1 (the ortholog of rice OSH1) and ZmFEA4 (ortholog of OsbZIP47). These together indicate likelihood of OsbZIP47 and OSH1 interactions for co-regulation of targets. In Arabidopsis, *pan ettin* phenotypes suggest *AtPAN* and *AtETTIN/ Auxin-Responsive Factor3* (*ARF3*) redundantly regulate floral organ numbers and patterns (Sessions et al., 1997). Additionally, rice *ETTIN1* and *ETTIN2* RNAi lines have aberrant plant height, compromised panicle branching and defects in stamen and carpel development (Khanday et al., 2013). Some of phenotypes resembled those observed in *OsbZIP47KD* plants. Thus, we hypothesized OsETTIN1 and OsETTIN2 may interact with OsbZIP47 to modulate aspects of organ development. Similarly, mutant alleles in *RFL,* the rice *AtLEAFY* ortholog, (*apo2* and *scc* allele) or *RNAi RFL* knockdown (Kyozuka et al., 1998; Rao et al., 2008; Wang et al., 2017) and mutants in *OSH1* (*osh1*) (Tsuda et al., 2009, 2011, 2014), the rice ortholog of *ZmKN1*, share some phenotypes of *OsbZIP47KD* plants. Common phenotypes are shorter plant height, delayed flowering, reduced panicle rachis length and branch complexity. Based on these meta-analyses, protein partnership between OsbZIP47 and ETTIN1, ETTIN2, RFL, and OSH1 was tested by the yeast two-hybrid (Y2H) assay. OsZIP47ΔC (lacking 186-385 amino acids from the C-ter was taken at bait protein in fusion with Gal4 BD and prey proteins (OsMADS1, or OSH1, or RFL, or ETTIN1, or ETTIN2) were fused to GAL4 AD. The GAL4AD-OsMADS15 interaction with GAL4DB-OsMADS1 was taken as the positive control (Moon et al., 1999; Lim et al., 2000). Also, homodimerization capability of OsbZIP47ΔC was tested. Based on growth pattern of transformed yeast cells on reporter media SD/-Leu-Ura-His + 5mM3AT and the X-gal quantitative assays we infer a strong interaction between OsbZIP47 and OsMADS1 and RFL whereas OSH1 and OsbZIP47 have moderate interaction (Fig. 5A and B). We also performed *in-planta* Bimolecular fluorescence complementation (BiFC) assays. *OsbZIP47ΔC* was cloned upstream to the coding sequence of C-terminal region of split YFP to express *OsbZIP47ΔC*-cYFP fusion protein. The CDS of *OsMADS1*, *OsbZIP47ΔC*, *RFL, OsETTIN1, OsETTIN2* and *OSH1* were cloned in frame downstream to coding sequence of the N-terminal split YFP (nYFP) to express nYFP fusion proteins. These six different combinations of nYFP and cYFP fusions were transiently co-expressed in onion epidermal cells. Nuclear YFP fluorescence signals confirmed protein interaction of OsbZIP47 with OsMADS1, OSH1 and RFL (Fig. 5C). Thus, we suggest that OsbZIP47 partnership with OsMADS1, OSH1, and RFL could contribute to meristem functions.

**Figure 5.**
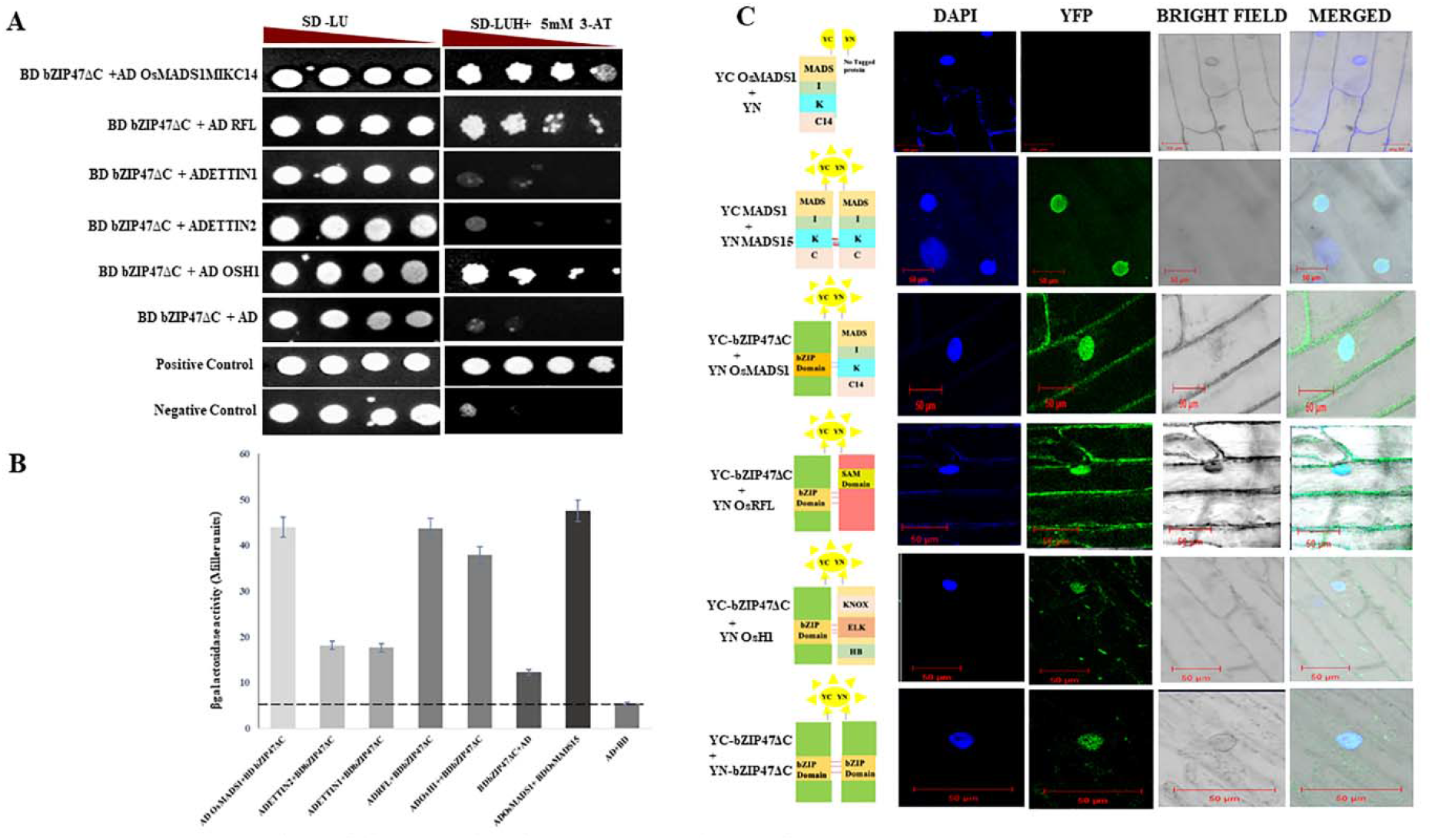
Interaction of OsbZIP47 with other meristem factors. (A)Yeast two hybrid (Y2H) assays with OsbZIP47ΔC protein (lacking 186-380 amino acids in C terminal) and predicted protein partners such as OsMADS1 (MIKC14 domain), RFL, OsETTIN1, OsETTIN2 and OSH1. Serial dilutions of yeast cells (PJ694A) spotted on media lacking histidine and supplemented with 5mM 3-AT. Growth after five days at 30 °C is shown. (B) Quantitative β-galactosidase activity assay of the indicated interaction pairs. ONPG was used as a substrate for detection of *lacZ* reporter gene expression levels and absorbance was determined at 420 nm. Yeast transformants with OsMADS15 in pGBDUC1 vector and OsMADS1 MIKC14 in pGADC1vector served as positive control for protein interaction. A combination of pGADC1 and pGBDUC1 empty vectors (without protein fusion) was the negative control. (C) Validation of yeast two hybrid protein interactions by Bimolecular fluorescence complementation (BiFC) assays in onion epidermal cells. YFP fluorescence was detected when OsbZIP47ΔC-cYFP fusion protein was co-expressed with OsMADS1-nYFP, RFL-nYFP, and OSH1 nYFP. Absence of YFP fluorescence in the negative control i.e., OsbZIP47ΔCYC and YN domain alone was the negative control. Bars=50 µm.

### Transcriptome of developing inflorescences of bZIP47 knockdown lines

To capture the gene expression landscape in the *OsbZIP47KD* panicles we carried out high throughput RNA-sequencing in two biological replicates of *OsbZIP47KD* and *WT* panicles (ln2-ln4, 1mm to 5mm panicles), and the differentially expressed genes (> two-fold change, p value <0.05) were extracted (Supplemental dataset S1; Supplemental Material and Methods). Among them, 1944 genes had lowered transcript levels in *OsbZIP47KD* hence during normal development their expression is likely upregulated by OsbZIP47. On the other hand, 856 genes had elevated transcript levels in knockdown tissues and thus these are the genes directly or indirectly downregulated by OsbZIP47 (Supplemental dataset S1). Gene Ontology (GO) analysis of these positively regulated gene sets (Fig. 6A and B; Supplemental Materials and Methods; Supplemental dataset S2) revealed that RNA (regulation of transcription), lipid and CHO metabolism, signalling, development and hormone metabolism were enriched in the positively regulated gene set (Fig. 6A). Whereas genes related to secondary metabolism, transport, cell wall, signalling, stress, hormone metabolism and miscellaneous were overrepresented in the negatively regulated set (Fig. 6B). Not surprisingly, genes involved in hormone metabolism are controlled by OsbZIP47 either positively and negatively (Fig. 6C and 6D). Specifically, Jasmonate (JA) and Abscisic acid (ABA) pathway genes are overrepresented in the positively regulated gene set. While Auxin, cytokinin (CK) and ethylene pathway genes are notable in the negatively regulated gene set, members of gibberellic acid (GA) pathway occur in both positively and negatively regulated gene sets (Fig. 6B). Here we give examples of *OsbZIP47* downstream genes that could interlink hormone pathways for panicle and floret development. OsAP2-39 balances the antagonistic relation between ABA and GA by modulating expression levels of *OsNCED1* (9-cis-epoxycarotenoid dioxygenase1) and *OsEUI1* (Elongated Upper most Internode1) for seed germination and plant development (Yaish et al., 2010; Shu et al., 2013, 2016). Interestingly, we observe positive regulation of *OsAP2-39* and negative regulation of *EUI1* by OsbZIP47. Thus, we suggest that OsbZIP47 enhances ABA biosynthesis and suppresses GA biosynthesis possibly to regulate plant height and rachis elongation (Fig. 2B and F). Also observed is reduced *SPL7* expression that regulates inflorescence meristem and spikelet transition (Dai et al., 2018) and reduced *OsMADS16* transcripts, a Class B stamen and lodicule organ identity gene (Nagasawa et al., 2003). These findings corelate well with the *OsbZIP47KD* inflorescence branching and floret organ defects (Fig. 2 and 3). Transcription factors control the dynamics of hormone signalling pathways by modulating gene expression levels. Among transcription factors genes that are deregulated in *OsbZIP47KD* panicles are – bHLH gene members (22), Co-like Zn finger (5 genes), TCP class 1 (2 genes) and TCP class 2 (1 gene), trihelix (4 genes), C2H2 zinc (13 genes), in the negatively regulated set (Fig. 6C, Supplemental dataset S2). The genes for transcription factors that are positively regulated by OsbZIP47 are from NAC class, WRKY class, and MYB-related class genes. Among the latter class are: *OsLHY* (*LATE ELONGATED HYPOCOTYL*) and *CCA1* (*CIRCADIAN CLOCK ASSOCIATED1* which work in a regulatory loop to control photoperiodic flowering (Lu et al., 2009) plant tillering and grain yield (Wang et al., 2020). We speculate that the delayed flowering phenotypes and reduced plant height of *OsbZIP47KD* plants can be associated with their positive regulation by *OsbZIP47*. The organogenesis regulator *YABBY* domain factor-*DROOPING LEAF* (*DL*) functions in both carpel specification and in leaf development (Yamaguchi et al., 2004) and the F-box gene, *APO1* with roles in spikelet and floret development are affected in *OsbZIP47KD* panicles (Supplemental dataset S1; Fig. 6E; Ikeda et al., 2005, 2007). The positive regulation of *CUC1* (*CUP-SHAPED COTYLEDON1*) by OsbZIP47 supports plausible mechanism for its influence on organ whorl boundaries (Takeda et al., 2011; Fig. 6E). We speculate that positive regulation of *DWARF AND RUNTISH SPIKELET2/FERONIA like Receptor 2* (*FLR2*) may contribute to architecture, fertility, and seed yield (Li et al., 2016; Fig. 6E). Overall, these results give a snapshot of OsbZIP47 molecular functions in inflorescence tissues and give leads to its unique *vs.* evolutionarily conserved roles for panicle meristem transitions, floral organ development.

**Figure 6.**
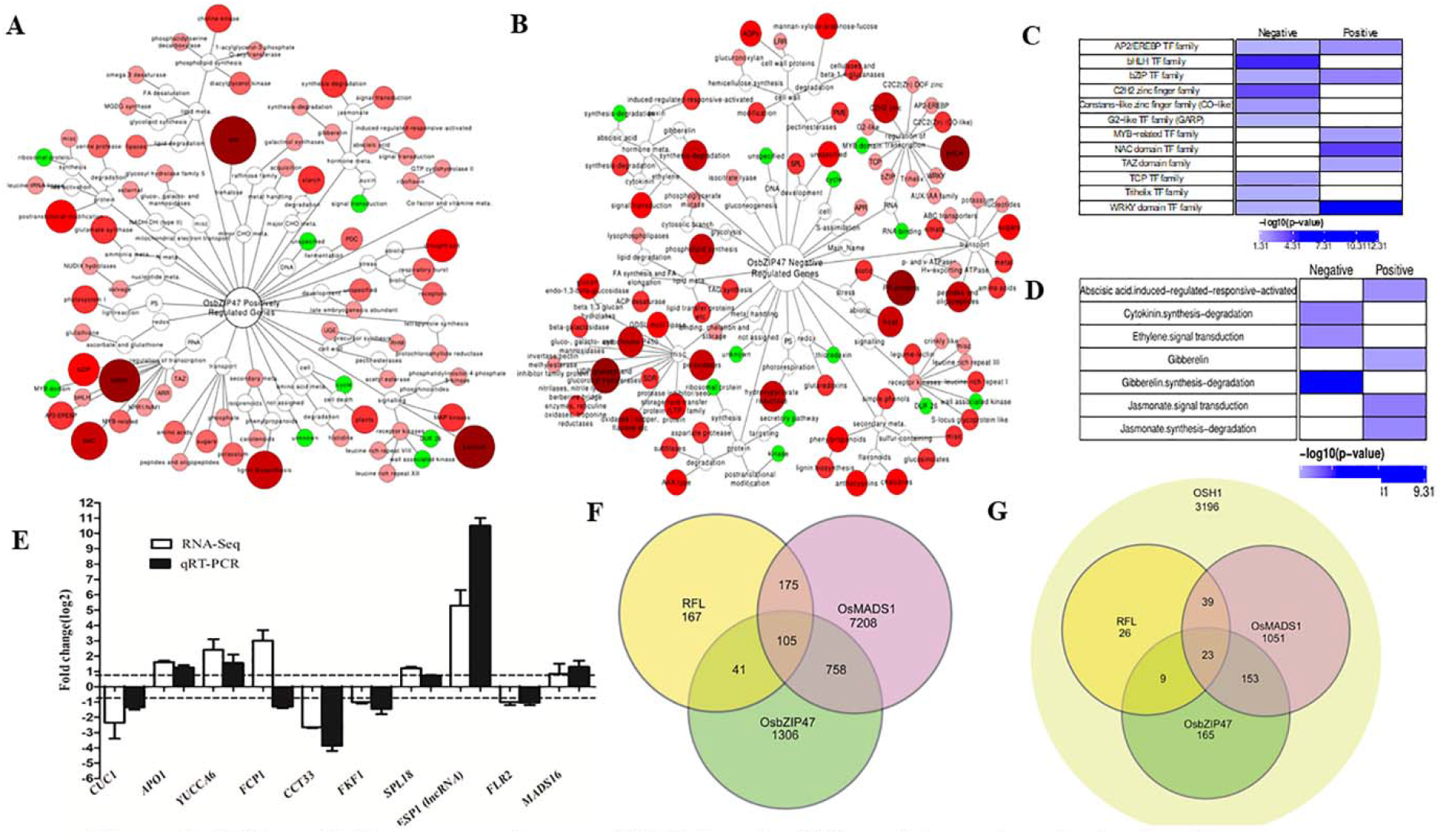
Differentially expressed genes (DEGs) and a GO enrichment analysis of pathways downstream to OsbZIP47. Enrichment networks depicting pathways regulated by OsbZIP47. (A) positively regulated, and (B) Negatively regulated pathways. The green and red color nodes represent enriched and depleted functional categories respectively (P <0.05). The size and color intensity of the node correlates with the over-representation of genes within the given class. (C) Enrichment map of transcription factor (TF) genes shows those encoding members from bHLH, G2-like, C2H2 zinc finger, TCP, Trihelix are underrepresented, whereas TFs such as AP2/EREBP, MYB-related, NAC domain, TAZ, WRKY, bZIP were overrepresented. (D) Enrichment analysis of genes for phytohormone metabolism and signaling. Cytokinin, Ethylene, Gibberellin pathways were enriched in negatively regulated dataset, whereas, Abscisic acid, Gibberellin, and Jasmonate pathways predominate in the positively regulated set. (E) RT-qPCR analysis of the relative fold change for transcripts from several candidate target genes of OsZIP47 in 0.1 to 0.5 cm panicles from *OsbZIP47KD* as compared to *WT* tissues. The normalization of transcript level of each gene was done using *UBQ5* transcriot levels. Error bars represents the standard deviation. (F) Venn diagram showing unique and overlapping sets of differentially expressed genes derived from *OsMADS1, OsbZIP47* and *RFL* transcriptome datasets (Rao et al. 2008; Khanday et al., 2013 and this study). (G) Genes implicated to be coregulated by *OsMADS1, OsbZIP47,* and *RFL* and bound by *OSH1* (Rao et al. 2008; Khanday et al., 2013; Tsuda et al., 2014 and this study).

### Comprehensive datamining of transcriptome datasets of OsMADS1, OsbZIP47, RFL and OSH1

Extending the findings of OsbZIP47 interaction with OsMADS1, OsbZIP47, RFL and OsH1, we carried out meta-analyses of published transcriptome datasets affected in mutants of these partner proteins. To identify candidate genes for co-regulation by these factors we examined the differential transcriptome in *dsRNAiOsbZIP47*KD, dsRNAi*OsMADS1* and dsRNAi*RFL* panicles (Khanday et al., 2013; Rao et al., 2008). First we aligned genes from three different transcriptomic datasets for this meta-study (Supplemental Materials and Method). 2210 deregulated genes in *dsRNAiOsbZIP47*KD panicles were examined for overlap with 8246 affected genes in *dsRNAiOsMADS1*KD transcriptome (Fig. 6F; Supplemental dataset S3; Khanday et al., 2013). 863 candidates for co-regulation by OsbZIP47 and OsMADS1 were deciphered (Fig. 6F Supplemental dataset S3); 204 of them were down-regulated in both *OsbZIP47KD* and *OsMADS1KD* lines. These include genes encoding *GASR3* (Gibberellin-regulated protein precursor expressed), *OsSAUR11, DWARF AND RUNTISH SPIKELET2* (*FLR2),* and *TIFY* (ZIM domain transcription factor). Another group of genes (386 out of 863 genes) were up-regulated in both *OsbZIP47KD* and *OsMADS1KD* panicle datasets. This sub-set consist of genes encoding for transcription factors that regulate hormone signalling such as, *OsIAA20, YUCCA7, PIN5B* (*PROBABLE AUXIN EFFLUX CARRIER COMPONENT 5B), EIL4*, (*ETHYLENE INSENSITIVE 3 LIKE 4*) gene, *GNP1* (*GRAIN NUMBER PER PANICLE1*) and *OsMADS16.* Among the 863 candidate genes co-regulated by OsbZIP47 and OsMADS1, is a subgroup of 153 genes (Fig. 6G) that are also bound by OSH1(Tsuda et al., 2014). Striking among this sub-set are: *IAA20*, *YUCCA7, OsMADS27, SPLIT-HULL* (*SPH*) and *DWARF AND RUNTISH SPIKELET2/FERONIA like Receptor 2* (*FLR2*). Similarly, 146 genes (Fig. 6F) are common to the differentially expressed genes in *OsbZIP47KD* panicles (RNA-Seq) and a low density microarray study of panicles from dsRNAi*RFL* knockdown plants (Rao et al., 2008). Interestingly, 33 out of 146 genes were down-regulated in both these datasets including ethylene signalling gene *ACO1* which regulates internode elongation (Iwamoto et al, 2010), and jasmonate signalling gene *TIFY11D* (Kim et al., 2009), and the *EMBRYOSAC1* (*OsEMSA1*), involved in embryo sac development (Zhu et al., 2017). Lastly we identify 105 differentially expressed genes common to the differential transcriptome in knockdown of *RFL*, *OsMADS1*, and *OsbZIP47* panicles (Fig. 6F) The notable genes include ethylene insensitive-like gene 2 (*EIL2*), allene oxide synthase (*AOS1*), and gibberellin 2-oxidase *9* (*GA2OX9*). A sub-set of 23 genes are potentially regulated by OSH1 (Fig. 6G, Tsuda et al., 2014). We infer that these transcription regulators possibly multimerise in one or more forms of complexes to regulate the meristem development in rice. Overall, these findings hint that protein complexes, plausibly heterogeneous, with combinations of OsbZIP47 and its varied partner factors may spatially and temporally co-ordinate downstream gene expression during panicle development, spikelet, and floret development.

### Redox dependent DNA binding of OsbZIP47

DNA binding by Arabidopsis AtPAN is redox-sensitive due to the five Cysteine residues in the extended N-terminal domain (Gutsche and Zachgo, 2016; Supplemental Fig. S8). Unlike PAN homologs from diverse species rice *OsbZIP47* lacks this domain. All proteins share a conserved Cys in the C terminal transcription transactivation domain (AtPAN Cys340/ OsbZIP47Cys269). Cys17 in OsbZIP47 is conserved in monocot species, while Cys196 is unique to *OsbZIP47.* Thus, though the rice and *Arabidopsis* proteins are close homologs they differ in size and in the overall number of cysteine residues. To examine OsbZIP47 oligomerisation and the effects of its redox status on binding to target gene *cis* DNA elements, the full length OsbZIP47 protein was expressed in bacteria. The DNA binding activity of AtPAN is regulated by S-glutathionylation of the conserved Cys340 by AtROXY1, a glutaredoxin redox enzyme (Li et al., 2009; Gutsche and Zachgo, 2016). The corresponding conserved Cys 269 in OsbZIP47 may also render the rice protein to be redox sensitive for biochemical activity. To determine if the protein forms higher order self-oligomers, size exclusion chromatography with the purified TrxHis OsbZIP47 (62Kda) protein was done and the elution of the protein in the column void volume (Fig. 7A) suggested either aggregation or the formation of high order oligomers. To examine the latter possibility the purified OsbZIP47-Trx protein was treated with 2mM diamide, an oxidizing agent. Another aliquot of the protein was in parallel treated with 20mM DTT, a reducing agent, and both treated protein fractions were analysed on a non-reducing SDS-PAGE gel. The oxidized OsbZIP47-His-Trx sample had slower mobility whereas the reduced bZIP47-His-Trx sample migrated with the expected mobility for a ∼68Kda protein. Importantly, we found that the effects of the oxidizing agent (diamide) can be reversed by DTT treatment. These data show OsbZIP47 oligomerisation is affected by its redox-status (Fig. 7B). The DNA binding of OsbZIP47 was tested at the TGACGT *cis* motif (predicted for bZIP DNA binding) at -371bp in the *OsFCP1* locus; a downstream gene target whose expression levels are affected in the SAM and in panicles of *OsbZIP47KD* plants (Fig. 7C; Fig. 1G). EMSA assays with full-length OsbZIP4762Kda protein (Fig. 7D) were compared with the partial 34.3Kda OsbZIP47-DNA-Binding Domain (DBD) alone (Fig. 7E). OsbZIP47 binding to DNA was redox dependent despite the absence of the extended Cys rich N-ter domain commonly found in homologs from other species.

**Figure 7.**
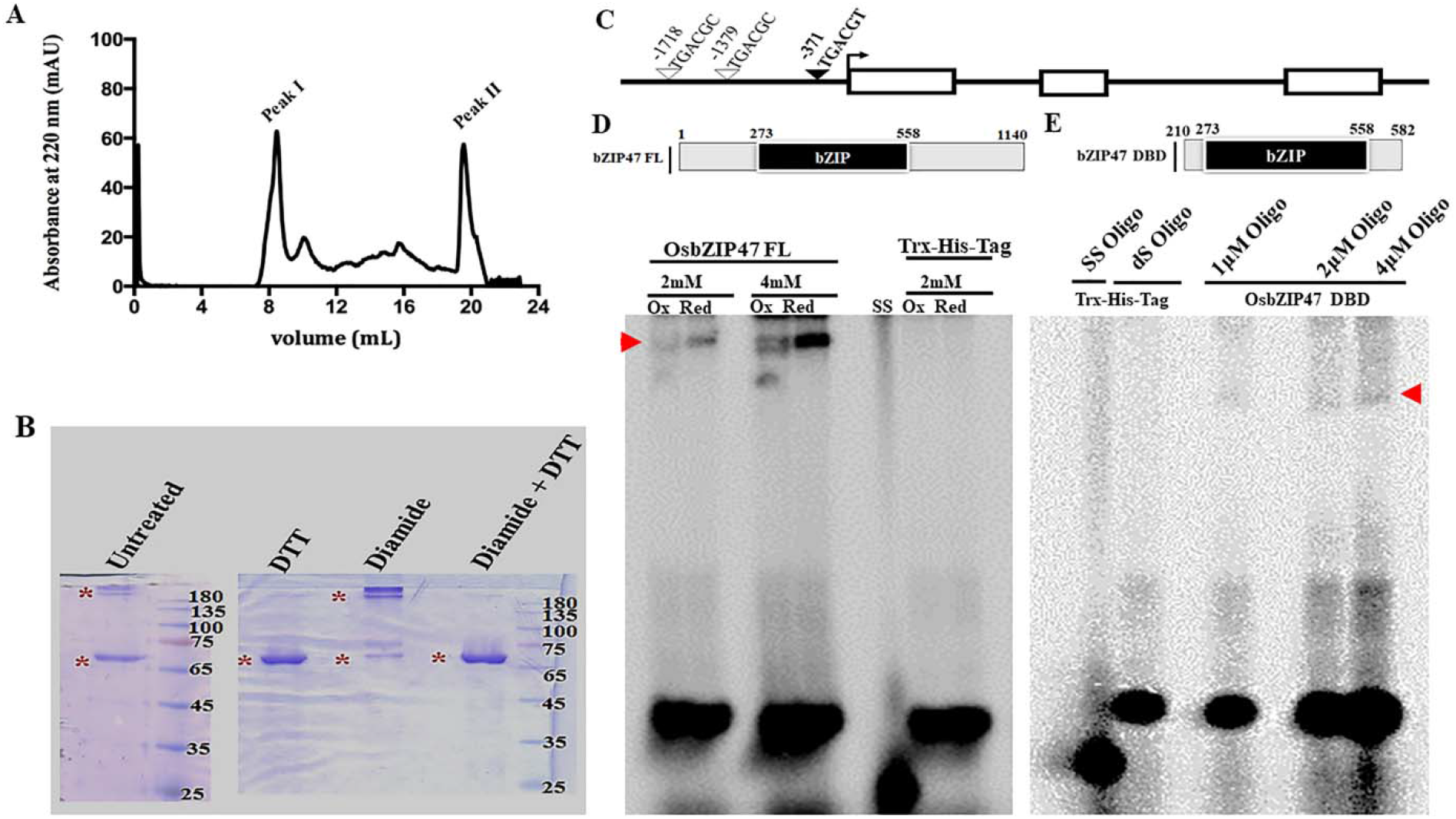
OsbZIP47 oligomerisation and the effects of its redox status on DNA binding. (A) Size-exclusion chromatography (SEC) profile for OsbZIP47 protein/oligomer on a Superdex 200 increase column. (B) Mobility of purified OsbZIP47-Trx protein on non-reducing 10% SDS-PAGE after treatment with 2mM diamide (an oxidizing agent), or with 20mM DTT (a reducing agent), or with 2mM diamide followed by 20mM DTT. (C) Schematic representation of the *FCP1* gene locus. Exons are open white boxes, introns are shown as black lines, and the predicted *cis* motifs for bZIP DNA binding domain (DBD) are represented as inverted triangles. The DNA binding properties of OsbZIP47FL and partial OsbZIP47DBD proteins was tested for the TGACGT *cis* motif located at -371bp of *FCP1* (marked with a filled inverted triangle). (D and E) EMSA assays with full-length OsbZIP47 (62Kda protein) as compared to the partial protein of 34.3Kda containing OsbZIP47 DBD. Retarded mobility of DNA-protein complexes (red arrow) for the *FCP1* target site is indicated. Unbound dsDNA oligonucleotide are at the bottom.

## DISCUSSION

The DNA binding domain in Arabidopsis PAN, maize FEA4 and rice OsbZIP47 proteins is conserved (Chuang et al., 1999; Nijhawan et al., 2008). Hence any species-specific developmental roles for these factors may arise from interacting co-regulators, from protein modification and through variations in downstream pathways that they regulate. Arabidopsis *AtPAN* has pleiotropic vegetative and reproductive growth effects. Flowers in the *pan* mutant are characterized by an increase in floral organ number, without a corresponding increase in FM size (Running and Meyerowitz, 1996; Chuang et al.,1999). Another study reported early flowering in long days for the *pan* mutant, while plants grown short-days had enlarged SAM and inflorescence meristems (Maier et al., 2011). Maize mutant *zmfea4* have enlarged vegetative SAMs, severely fasciated inflorescences and florets with reduced stamen numbers (Pautler et al., 2015). We show *OsbZIP47KD* plants have abnormalities in shoot meristem size homeostasis that are similar to maize *fea4* mutant. Yet other phenotypes are unique to rice *OsbZIP47KD* lines for instance: delayed flowering, increased stamen numbers, chimeric floral organs and subtle changes to grain size and shape. The partnership of OsbZIP47, with meristem regulators-OsMADS1, RFL and OSH1 (KNOX1/ STM), its oligomeric and redox status could relate to its functions in different meristems. These partnerships and the findings that emerge from the differential transcriptome in *OsbZIP47KD* panicles allowed us to map downstream genes regulated by OsbZIP47, and those potentially dependent on its co-regulators.

### Meristem maintenance and development in vegetative and reproductive phase

In *OsbZIP47KD* plants the variability in shoot meristem size, reduced size in 5 DAG seedlings while enlarged in 25 DAG seedlings, hints at the failure of compensation mechanisms that maintain meristem size. The enlarged SAM in 25 DAG *OsbZIP47KD* seedlings is superficially similar to shoot meristems of maize *fea4*, yet there are underlying subtle differences in rice knockdown plants. A detailed analysis of cell size in the SAM of *OsbZIP47KD* seedlings showed unexpected increased size in L1 cells and those underlying it that co-relates with altered SAM size. The meristem functions of Arabidopsis PAN and Maize FEA4 are independent of or are in parallel to the core CLV-WUS pathway that controls meristem size (Running and Meyerowitz 1996; Running et al. 1998; Paulter et al., 2015). The reduced expression levels of rice *FCP1,* a *CLV3*-like gene with functions in vegetative SAM homeostasis, downregulation of *CUC1, APO1, YABBY6, CYP734A4* gene expression suggest several complex pathways by which OsbZIP47 contributes to vegetative shoot meristems and lateral organ (leaf) differentiation. Interestingly our meta-analysis of genes potentially co-regulated by OsbZIP47 and its partners (Fig. 6) Tsuda et al., 2014) point to *CYP734A*4 as one of the common gene targets controlled by OSH1-OsbZIP47 protein partnership. The down regulation in expression levels of this factor for BR biosynthesis in *OsZIP47KD* tissues and certain meristem phenotypes in 25 DAG meristems of *OsZIP47KD* plants and in *cyp734a4* mutant (Tsuda et al., 2014), supports a mechanism by which this partnership influences meristem and lateral primordia development. Cytokinin (CK) in conjunction with other phytohormones, auxin and Gibberellic acid is essential for cell division and organ differentiation (Leibfried et al., 2005 and Zhao et al., 2010; Su et al., 2011). Transcriptome profiling of *OsbZIP47KD* panicles shows deregulated expression levels for *Knotted1-like11*, *IPT1*, *IPT2*, *IPT8*, *GA3Ox3*, *GA2Ox4, GA20Ox3*, *YUCCA6* and *YUCCA7* and as well for the rice CLV-peptide family genes *FON2* and *FCP1.* We propose OsbZIP47 functions as an integrator of the WUS-CLV and WUS-KNOX pathways for meristem development as it regulates expression levels of key factors in both pathways. Also, genes predicted to have roles floral organ primordia differentiation genes such as *OsBLH1* (BEL1-like homeodomain), *OsKANADI* and *CUC1* are deregulated in panicle tissue of *OsbZIP47KD*. Interestingly, vegetative meristem size and reproductive panicle branching phenotypes of *OsbZIP47KD* panicles resemble *apo1* and *apo2/rfl* mutants (Ikeda et al., 2005, 2007, 2009; Rao et al. 2008; Deshpande et al., 2015). Thus, the interactions between RFL and OsbZIP47 could positively regulate the panicle meristem branch identity and its developmental transitions. A significant example of a candidate target gene for co-regulation by ObZIP47 and RFL is *CUC1*. Further, the elevated transcript levels for *APO1* in *OsbZIP47KD* panicle tissues hints that *OsbZIP47* in the wild type panicle supresses the expression of *APO1* which we speculate affects its partnership with *APO2/RFL.* With this we anticipate OsbZIP47 could have evolved to regulate of some unique molecular pathways for vegetative and reproductive phase meristem growth and development.

### Transition of Shoot apical meristem (SAM) to Inflorescence meristem (IM)

Unlike the early flowering in long days seen in the Arabidopsis *pan* mutant, knockdown of rice *OsbZIP47* showed delayed flowering. Importantly, the latter is a common trait in mutants or knockdown transgenics in *OsMADS1, OSH1* and *APO2/RFL* that encode OsbZIP47 protein partners (Jeon et al., 2000; Rao et al., 2008; Tsuda et al., 2011; Ikeda-Kawkatsu et al., 2004; Fig. 2). These observations support our hypothesis of functions for these factors in “one or more” complexes. Panicle transcript analysis in knockdown transgenics indicates OsbZIP47 can promote flowering by fine-tuning the expression of several flowering time regulators that are upstream to florigens *Hd3a* and *RFT* (e.g., *OsLF*, *OsLBD38*, *OsLBD38, OsIDD6*) and by controlling expression of circadian clock-associated genes like *OsFKF1*, *OsPCL1* and *OsLHY/CCA1*. Several rice flowering time QTLs also influence grain traits (Zhu et al., 2017; Chen et al., 2014; Ma et al, 2019). The subtle effects of *OsbZIP47KD* on rice grain shape, are not reported for maize *fea4* kernels suggesting unique effects in rice grain size and shape. Our transcriptomic analysis identified a set grain shape genes: *OsLG3* (*Long Grain 3*), *GS9* (*Grain Shape Gene on Chromosome 9*), *GW7* (*Grain width QTL on chromosome 7*), *FLO2* (*Floury Endosperm 2*) as genes plausibly regulated by OsbZIP47. *GS9* positively controls the grain size by altering cell division along with BR signaling (Zhao et al., 2018). Interestingly, we noted high level expression of Cyclin D (*CYCD7*) in *OsbZIP47KD* panicles (Supplementary dataset S1). This is noteworthy as in Arabidopsis the tissue and stage specific control of this G1-S phase cell cycle gene controls cell division in different contexts, with ectopic expression driving abnormal cell division patterns (Weimer et al., 2018). Additionally another study showed tight control of cell division is important during Arabidopsis seed development as overexpression of *CYCD7*, which is not normally expressed in seeds, increased cell division and expansion in embryo and the endosperm (Collins el al., 2012). With this we postulate that *OsbZIP47* links flowering time, cell cycle and brassinosteroid signaling to regulate grain shape.

### Regulation of inner floral organ identity and specification

Consistent with *OsbZIP47* expression domain in near mature florets (Sp6-Sp8), we observed lodicule and stamen differentiation defects in *dsRNAiOsbZIP47* florets. Increased stamen numbers, with degenerated anthers on short filaments and the partial homeotic transformation of stamens to lodicules support roles for OsbZIP47 in organ differentiation. Interestingly, in *OsbZIP47KD* florets the higher transcript abundances of *OsMADS16* (homolog of *AtAP3*) and *DL*, a Class C function contributor in rice (Supplemental dataset S2), are indicative of some distinct effects in rice florets. Overexpression of *OsMADS16* can increased stamen numbers and form stamenoid carpels without any effects on lodicules (Lee et al., 2003). More recently, rice transgenics with a modified repressive *OsMADS16* (OsMADS16-SDX repressor domain fusion) generated indehiscent anthers (Sato et al., 2012). These phenotypes are akin to third whorl organ differentiation defects seen in *OsbZIP47KD* florets. Microsporogenesis in anthers of Arabidopsis flowers requires *SPOROCYTELESS/NOZZLE* (*SPL/NZZ*) that is a gene target of Class, B and C organ identity factors (Ito et al., 2004). In line with these reports, we noted upregulation of *OsSPL*, possibly an effect of increased *OsMADS16* in *OsbZIP47KD* florets. Therefore, we conclude *OsbZIP47* regulates stamen differentiation by regulating Class B and *OsDL* a rice specific Class C function contributor. *FON2* and *APO1* genetically interact with *OsMADS16;* further *OsDL* and *FON2* positively regulate *OsMADS16* expression (Ikeda et al., 2007; Xu et al., 2017). In *fon2* and *apo1* loss-of-function mutants abnormal lodicules and carpels with increased number of stamens are known; whereas in their gain-of-function results in reduction of all organs corelated with reduced floret meristem size (Suzaki et al., 2006; Ikeda et al., 2009). On, comparing *OsbZIP47KD* phenotypes to those in *apo1* or *fon2*, we note opposing affects, where *FON2* and *APO1* expression levels are increased. Therefore, we propose that *OsbZIP47KD* floral stamen phenotypes are *OsMADS16* mediated and likely independent of *FON2* and perhaps is redundant with *APO1* (Ikeda et al., 2007; Xu et al., 2017).

### Biochemical properties of OsbZIP47 can underlie its unique functions and downstream effects

Multiple sequence alignment shows OsbZIP47 amino acid sequences share 50% or greater identity with homologs across diverse species. A common feature among many bZIP47 proteins is a variable extended N terminal domain, except in OsbZIP47 and Bamboo PH01000727G0540 (Supplementary Fig. S8). DNA binding activity of Arabidopsis PAN is a redox-sensitive binding due to the presence of five Cysteine (Cys) amino acids in the extended N-ter domain and the conserved C-ter Cys340 in the transcription transactivation domain (Gutsche and Zachgo, 2016). In contrast, the shorter OsbZIP47 protein has three Cysteines (Cys17, Cys196 and Cys269). Cysteine17 of OsbZIP47 is represented in all the monocot species, Cys 269 of OsbZIP47 (Cys 340 of AtPAN) is conserved in all plant homologs compared here (Supplemental Fig. 8), whereas Cys 196 is unique to OsbZIP47. Interestingly, among the proteins compared here wheat TAE56722G002 has the maximum number of 11 Cysteines. These observations hint that the number of Cys residues in this clade of bZIP proteins may have evolved for species-specific roles plausibly for the adoption of unique structures with effects on tissue-specific target gene expression. Interestingly, OsbZIP47 showed redox-dependent DNA binding despite being a short protein with reduced number of Cysteines residues. In rice OsGRX19 or MICROSPORELESS1 (OsMIL1) is the potential glutaredoxin redox enzyme for OsbZIP47 as it is the homolog of the glutaredoxin redox enzyme AtROXY1 and ZmMSCA1 from Arabidopsis and Maize (Timofejeva et al., 2013; Yang et al., 2015). Indehiscent anthers is a phenotype of common to *OsbZIP47KD* and *mil1* mutant (Hong et al., 2012) leading us to propose S-glutathionylation of OsbZIP47 could be important for anther development.

Overall, we uncover conserved as well as unique roles and mechanisms of OsbZIP47 that support meristem maintenance and determinacy, in vegetative growth and reproductive development leading to grain formation. These functions make OsbZIP47 a potential locus for a potential locus for allele mining and crop improvement.

## Materials and Methods

### Plasmid Constructs Generation and Rice Transformation

For siRNA (interference) mediated knockdown of endogenous *OsbZIP47*, a gene-specific 226bp 3’UTR DNA fragment was cloned in the sense and in the antisense orientation in same vector (pBluescript) and were separated by a 270-bp linker fragment. Subsequently, this insert was subcloned under maize ubiquitin promoter in the binary rice expression vector pUN for expression of *OsbZIP47* hairpin RNAs (Supplementary Figure S1; Prasad et al., 2001). For generating over-expression of *OsbZIP47* the full length cDNA was cloned at BamHI (blunted)-KpnI site in pUN vector to yield recombinant *pUbi*::*OsbZIP47* (Supplementary Figure S5). These constructs for knockdown and over-expression of *OsbZIP47* were transformed into *Agrobacterium tumefaciens* strain LB4404 and then co-cultivated with embryogenic calli from TP309 WT (*Oryza Sativa* var *japonica*) seeds as described by Prasad et al. (2001). Transgenic plants, dsRNAi *OsbZIP47* and Ox-*OsbZIP47*, were grown in IISc, Bangalore, Green house condition, ∼27°C during the months of January-May and July-October each time with control wild type plants.

### Phenotypic characterization of knockdown transgenic

dsRNAi *OsbZIP47* and Ox-*OsbZIP47* transgenic plants (T1, T2 or T3) were selected on half-strength MS medium containing 50 mg L^−1^ hygromycin. Detailed phenotypic analysis of dsRNAi *OsbZIP47* transgenic was performed in the T3 generation and for Ox-*OsbZIP47* transgenics in T1 plants. Tissue sections (7μm, Lecia microtome; RM2045) of 5 and 25 days seedlings were stained with Eosin-Haematoxylin staining and imaged by Apotome2 Zeiss. The size of cells in the SAM was measured using ImageJ. The seedling height of dsRNAi*OsbZIP47* KD lines was measured 7 day after germination (DAG). Adult plant height, lamina joint angle, panicle length, panicle branch characteristics, spikelet numbers were measured after panicle booting. Pre-anthesis florets, from panicles prior to their emergence from the flag leaf, were imaged using Leica Wild M3Z stereomicroscope.

### RNA-Sequencing and RT-qPCR

Next generation Sequencing (NGS) of RNA from *OsbZIP47* KD panicles (0.1cm-0.5cm) was done for two biological replicates with matched WT panicles tissues as controls. The total RNA was extracted using Trizol Reagent (Sigma) according to manufacturer instructions. 1μg total RNA was used for library preparation using rRNA depletion-based NEB Next UltraII RNA kit. NGS was performed using Illumina Hi-Seq, pair-end 2X150bp chemistry. After quality check (using FastQC and multiQC software) the reads were mapped against indexed *Oryza sativa* ssp. *japonica* cv. reference genome (RAP-DB; https://rapdb.dna.affrc.go.jp/) using STAR2 (v2.5.3a). Further differential gene expression (DGE) of read counts between wild type and transgenic were computed using edgeR (v3.28.0) package with the absolute log2 fold change ≥ 1 and ≤ −1 with p-value ≤ 0.05. For real-time qPCR experiments oligo(dT)-primed cDNAs were synthesized using 2ug of total RNA with MMLV (reverse transcriptase, NEB). qRT-PCR reactions were set up with 50-70ng of cDNA, 250 nM gene-specific primers and FastStart Universal Sybr Green Master (Rox) mix (Roche) in CFX384 real-time system (Biorad) or Applied Biosystems ViiA 7 system. Fold change in transcript levels for deregulated genes was calculated as difference in cycle threshold value between knockdown transgenic and wild type. To obtain normalized threshold value (ΔΔCt) first ΔCt value was calculated by subtracting the Ct value for internal control, *Ubiquitin5* from the Ct value for each gene of interest (Gene Ct-Ubi5 Ct). Then ΔΔCt was calculated by subtracting the wild tyope ΔCt value from the ΔCt value obtained for transgenic tissue. The fold change was calculated as 2^^-(ΔΔCt)^. Primers used and their sequences are given in Supplemental Table S2.

### RNA In Situ Hybridization

To generate *OsbZIP47* riboprobes, a gene-specific 226bp DNA fragment from 3’UTR (1329-1555 bp) was PCR amplified and cloned in the pBluescript KS+ vector. Sense and antisense Digoxigenin-labeled (DIG-UTP, Roche) riboprobes were prepared by *invitro* transcription using T3 and T7 RNA polymerases (NEB), respectively. Tissue processing and probe hybridizations was done in Prasad et al. (2005). Signal was developed using anti-digoxygenin-alkaline phosphatase (AP) conjugated antibodies (Roche) and BCIP (5-Bromo-4-chloro-3-indolyl phosphate)-NBT (nitro blue tetrazolium) chromogenic substrates (Roche). Images were captured by Apotome2 Zeiss microscope system.

### Bacterial Expression of OsbZIP47 FL protein and studies of oligomeric status

For OsbZIP47 protein expression and purification from bacteria, *OsbZIP47* full-length (FL) CDS and *OsbZIP47* DBD (240-582bp) was first cloned in the pET32a vector. Thioredoxin-His-tagged OsbZIP47FL and OsbZIP47DBD proteins were expressed from Rosetta (DE3) bacterial strain induced with 0.2 mM IPTG (Isopropyl β-d-1-thiogalactopyranoside) for 3 h at 37°C. Oligomeric states of OsbZIP47 protein was determined by analytical size exclusion chromatography (SEC) performed at 4°C on a Superdex 200 increase column (GE Healthcare *Inc*.) pre-equilibrated with buffer (25 mM sodium phosphate (pH 7.4) 100 mM NaCl and 5% glycerol). Approximately 400 μg protein, (∼2mg/ml) was injected into AKTA purifiers (GE healthcare *Inc.*) connected to the column. The flow rate was maintained at 0.3 ml/min and the protein elution profile was at 220 nm. The molecular weight was calculated using a standard plot. Molecular weight was calculated using the equation: Y= −0.602X + 4.6036, where, Y= V_e_/V_o_ (V_e_ =Elution volume; V_o_ = Void volume) and X= Log of molecular weight in Dalton.

### Electrophoretic mobility shift assays (EMSA)

*E coli* rosetta (DE3) bacterial lysates with the Trx-His-OsbZIP47 FL and OsbZIP47-DBD proteins were prepared in the buffer: 10mM HEPES-KOH, pH 7.8, 50mM NaCl, 0.5% Nonidet P-40, 0.5 mM EDTA, 1 mM MgCl2, 10%glycerol, 0.5 mM DTT, and 1x protease inhibitors cocktail (Sigma). 1-4μl lysate was incubated with 5’end P^32^ labelled DNA oligonucleotide probes for 30 minutes at 4°C in 1x EMSA buffer (20mM HEPES-KOH pH 7.8, 100 mM KCl, 2mM DTT, 1mM ETDA, 0.1% BSA, 10ng Herring sperm DNA, 10% Glycerol, 1X Protease Inhibitor cocktail) in a 15μl reaction volume. The binding reaction was resolved on a 8% native-PAGE gel in 0.5x Tris-borate EDTA (TEB) buffer at room temperature. The gel was analyzed by autoradiography in a phoshorimager (GE; Typhoon FLA 9500). The sequences of the EMSA probes are listed in Supplemental Table S2.

### Yeast Two-Hybrid assays

The full length CDS of *OsbZIP47* was amplified from KOME clone AK109719 using gene-specific primers and cloned into pBSKS vector. After verification by restriction digestions and sequencing, it was subsequently cloned into yeast two hybrid vectors, pGBDUC1 and pGADC1. Similarly, all the CDS fragments that would encode prey proteins such as OsMADS1, OsETTIN1/2, RFL, OSH1 and OsMADS15 were PCR amplified from either KOME cDNA clones or from cDNA made from panicle tissue RNA, and sub-cloned into destination pGBDUC1and pGADC-1 Y2H vectors. The bait clone pGBDUC1OsbZIP47 and indicated prey clones in pGADC1 vector were co-transformed into the yeast pJ69-4A Y2H strain (James et al., 1996) using the lithium acetate (LiAc) method. Transformants were selected on synthetic drop out media lacking Leucine and Uracil. Protein interactions were assessed in at least five purified transformants by serial dilution spotting of broth cultures onto SD/-Leu-Ura-His plates supplemented with 5mM of 3AT and by the ONPG assay (Supplemental Material and methods).

### Bimolecular fluorescence complementation assays (BiFC)

*OsbZIP47* cDNA with a truncated C domain (amino acid 199-385) was cloned into pSPYCE (M) (C-terminal fusion) and pSPYNE (R) 173 (N-terminal fusion) BiFC vectors (Waadt et al., 2008). Similarly, the full-length CDS encoding prey proteins such as OsMADS1, OsETTIN1/2, RFL, OSH1 and OsMADS15 were subcloned into pSPYNE (R) 173 vector. Six combinations of cEYFP and nEYFP fusions, including positive and negative controls, were transiently co-expressed in onion (*Allium cepa*) epidermal cells by *A. tumefaciens* (C58C1) infiltration as described in Xu et al. (2014). Co-transformed tissues were incubated at 25°C in dark for 48 h before being assayed for YFP activity. Fluorescence images were screened using a confocal laser microscope (Zeiss LSM880, Airyscan) with 2AU 480nm excitation and 520nm emission for detection of YFP signal.

### Meta-analysis

The published transcriptome datasets in dsRNAi*OsMADS1* and dsRNAi*RFL* panicles were adopted in this study to compare with that of *OsbZIP47KD* transcriptome dataset. The differentially expressed genes from each dataset was taken up for pair wise comparison to identify unique, or commonly (up-regulated, or down-regulated) downstream genes. The deregulated genes were also corelated with the published data on OSH1 genome-wide binding (Supplemental Material and methods).

## SUPPLEMENTAL DATA

**Supplemental Figure S1.** Schematic of T-DNA segment in the *pUbi:OsbZIP47*-RNAi construct and assessment of knockdown of *OsbZIP47* transcripts in transgenic line.

**Supplemental Figure S2.** Comparison of cell number in the L1 of 5 and 25 day old vegetative SAM tissues from in wild type *vs. OsbZIP47KD* seedlings.

**Supplemental Figure S3.** Spatial expression patterns of *H4*-Histone transcripts in 5 and 25-day SAM of wild type and *OsbZIP47KD* seedlings.

**Supplemental Figure S4.** Floral phenotypes of *OsbZIP47 KD* plants.

**Supplemental Figure S5.** Generation of *OsbZIP47* overexpression lines and phenotype in T1 generation plants.

**Supplemental Figure S6.** Comparative spatial expression of *OsbZIP47* with *OSH1* in panicles, spikelets and florets.

**Supplemental Figure S7.** Volcano plot and Gene Ontology pathway enrichment analysis of *OsbZIP4KD* transcriptome data.

**Supplemental Figure S8.** Multiple Sequence Alignment of OsbZIP47 with its orthologues showing conserved and non-conserved amino acid residues.

**Supplemental Table S1.** Quantifying phenotypes of *OsbZIP47* knockdown line #14 in the T3 generation

**Supplemental Table S2.** List of primers/Oligonucleotides used in this study.

**Supplemental Data Set S1**. List of genes deregulated by 2-fold (P < 0.05) in *OsbZIP47KD* line as compared to the wild type transcriptome.

**Supplemental Data Set S2.** List of DEGs for Gene Ontology Enrichment Analysis. **Supplemental Data Set S3.** List of genes co-regulated by *OsbZIP47, OsMADS1* and *RFL,* bound/unbound by *OSH1*.

## ACKNOWLEDGEMENTS

This work was funded by the Department of Biotechnology, Ministry of Science & Technology, Government of India Project entitled “Functional Analysis of Gene Regulatory Networks during Flower and Seed development in rice”, Project number: BT/AB/FG-1 (PH-II)/2009 to U.V.R. Funding to RR was from the UGC DSK fellowship: Project ID:No.F.4-2/2006 (BSR)/BL/1718/0151) and student fellowship to RP was from Indian Institute of Science. The infrastructure support for greenhouse facilities, confocal microscopy and phosphorimager was from DBT-IISc partnership program Phase II and this is gratefully acknowledged. Inputs of Dr. Grace Chongloi and from members of the UVR laboratory during the course of the study are also acknowledged. Ms Divya is thanked for assistance with confocal imaging and Mr Murthy, Mr Jagadeesh are thanked for assistance in plant growth and care.

